# Causal Role of the Medial Prefrontal Cortex in Learning Social Hierarchy

**DOI:** 10.1101/2023.11.23.568266

**Authors:** Chen Qu, Yulong Huang, Rémi Philippe, Shenggang Cai, Edmund Derrington, Frédéric Moisan, Mengke Shi, Jean-Claude Dreher

## Abstract

Social hierarchy is a fundamental principle of social organization and an important attribute of community stability and development. Yet, little is known about the causal role of specific brain regions in learning hierarchies. Here, using transcranial direct current stimulation (tDCS), we investigated the causal role of the medial prefrontal cortex (mPFC) in learning social and non-social hierarchies. In the Training phase, participants(N=128) acquired knowledge of social and non-social hierarchy in parallel, by trial and error. During the Test phase, they were presented with two items from hierarchies that were never encountered together and required to make transitive inferences. Anodal stimulation over mPFC impaired social hierarchy learning compared with non-social learning and this modulation was influenced by the relative social rank of the members (i.e. higher or lower status). Anodal stimulation also impaired transitive inference making, but only during early blocks before learning was established. Together, our results provide causal evidence of mPFC engagement in learning social ranks by observation.

## Main Text

We live in a social environment that is regulated by a variety of hierarchical structures ^1^. Optimization of our social interactions requires us to perceive status cues and continuously update hierarchical relationships, by determining the power of others relative to ourselves, to make social judgments in daily life ^2^. Social hierarchy is a group structure that exists in many species including non-human primates^3^, rodents^4^, fish^5,6^, and humans ^7,8^, which is crucial to maintaining the stability of populations and the health of individuals ^2,9,10^. Animal studies have shown that burtoni fish can infer the social hierarchy of competitors by observation learning ^6^, and clownfish can adjust their size and growth rate according to their hierarchical position in group ^5^. Similarly, social hierarchy affects human behaviors ^11^, such as decision-making ^12^ and empathy ^13^, enabling people to choose favorable alliances in social competition and avoid potential conflicts. Impairment in accurately monitoring one’s position in the social hierarchy can affect human health and increase the possibility of mental diseases such as social anxiety^10,14^.

Individuals can assess hierarchy information in several ways ^2^, including via the perception of dominance-related cues (faces with dominant features, body postures, etc.), observational learning by trial and error ^15,16^, and through competitive interactions ^17^. Although assessing the strength of competitors by dominance cues, such as body postures, facial expressions and physical attributes ^18–20^ is rapid, the information from such cues does not always coincide with the real hierarchy status. In contrast, learning dominant relationships through direct dyadic competitive interactions, by experiencing successive victories or defeats against competitors, is time-consuming ^17^, and may be costly in terms of potential physical injuries. Thus, learning social hierarchy by observation is an efficient way to acquire social hierarchy knowledge, without the cost incurred through competitive interactions.

Previous correlational fMRI studies indicated that learning social hierarchy engages the medial prefrontal cortex (mPFC), as well as the hippocampus ^15–17^. Using model-based functional magnetic resonance imaging (fMRI), Kumaran et al. (2016) developed an observational hierarchy learning task that distinguished training and test phases to study the neural representations of the process of acquiring knowledge of hierarchies (during a training phase) and making transitive inferences on the basis of that knowledge (during a test phase). The mPFC was involved when learning the power information about other individuals in social hierarchy ^21^. In contrast, the hippocampus was involved in general hierarchy learning, i.e., the learning of both Social and Non-social hierarchies ^16^. Recent models and experimental studies have proposed that the same brain representations that map space may be extended to a broad range of non-spatial problems in abstract cognitive space ^22–26^. These studies support that the mPFC and the hippocampus are involved in non-spatial relational memory tasks allowing them to make transitive inferences ^26–28^. Moreover, mPFC distinguishes between ranks higher and lower than oneself, and specifically shows reduced activity for trials involving higher social ranks^15^. The social comparison theory posits that people are driven to compare themselves with others for accurate self-evaluations ^29^. Thus, people compare themselves to others in two opposite directions which may differ in motivation and comparison target, etc. ^30,31^.

It remains unclear whether the mPFC, a key component of this mPFC-hippocampus network is causally necessary for two distinct processes needed to organize abstract relational information into a cognitive map: acquiring knowledge of the relative rank between two adjacent items and making transitive inferences between items never presented together before, to guide novel inferences. It is also unknown whether mPFC perturbation affects knowledge acquisition and/or the making of transitive inference processes in similar ways across the social vs non-social domains. More specifically, mPFC perturbation may have distinct effects depending on whether the knowledge is pertinent to higher or lower social ranks.

Here, we applied transcranial direct current stimulation (tDCS) to explore the causal relationship between the mPFC and hierarchy learning processes. tDCS is a noninvasive brain stimulation method that modulates neural excitability of specific brain regions using a low electrical current ^32^. Our aims were thus: (i) to investigate whether the mPFC plays a causal role when learning hierarchies or during the transitive inference processes; (ii) to establish whether the mPFC plays a causal role only for learning social but not for non-social hierarchy and whether this is influenced by relative social rank (i.e. higher or lower status).

Using a double-blind sham-control, and online stimulation design, participants were randomly assigned to receive either anodal (n=42), cathodal (n=42), or sham (n=44) stimulation over the mPFC (see Fig. 1A&B). A stimulation montage Fpz-Oz with 1.5mA current was adopted, using EEG10-20 system for electrode placement, across subjects (see Methods). As illustrated in Fig. 1C, the electric field simulation shows that the voltage gradient spread through the prefrontal cortex and targeted mPFC (MNI: -6, 46, 12; from Kumaran et al. 2016; see Methods). During brain stimulation, participants performed a Hierarchy Learning task (see Fig. 2D-F), including Training and Test phases for both social and non-social conditions. Self-information was added to the social condition to study how social ranks modulate the self-other comparison process of hierarchy learning. In the Training phase, participants were required to view pairs of hierarchically adjacent pictures and indicated which picture they thought had a higher status/power (Social) or more minerals (Non-social), with the correct feedback for each trial. Thus, through trial and error, they can update the hierarchical knowledges. In the Test phase, they were required to use the hierarchy information acquired during the Training phase to make transitive judgments concerning the hierarchical relationship between two non-adjacent entities (i.e, that were never seen together during training), with no feedback provided. They were also required to rate their confidence level from 1 (guess) to 3 (very sure), which allowed us to track the uncertainty of participants choices during the hierarchy learning. The training and test phases allowed us to investigate hierarchy knowledge updating (Training) and transitive inferences (Test).

**Figure 1.**
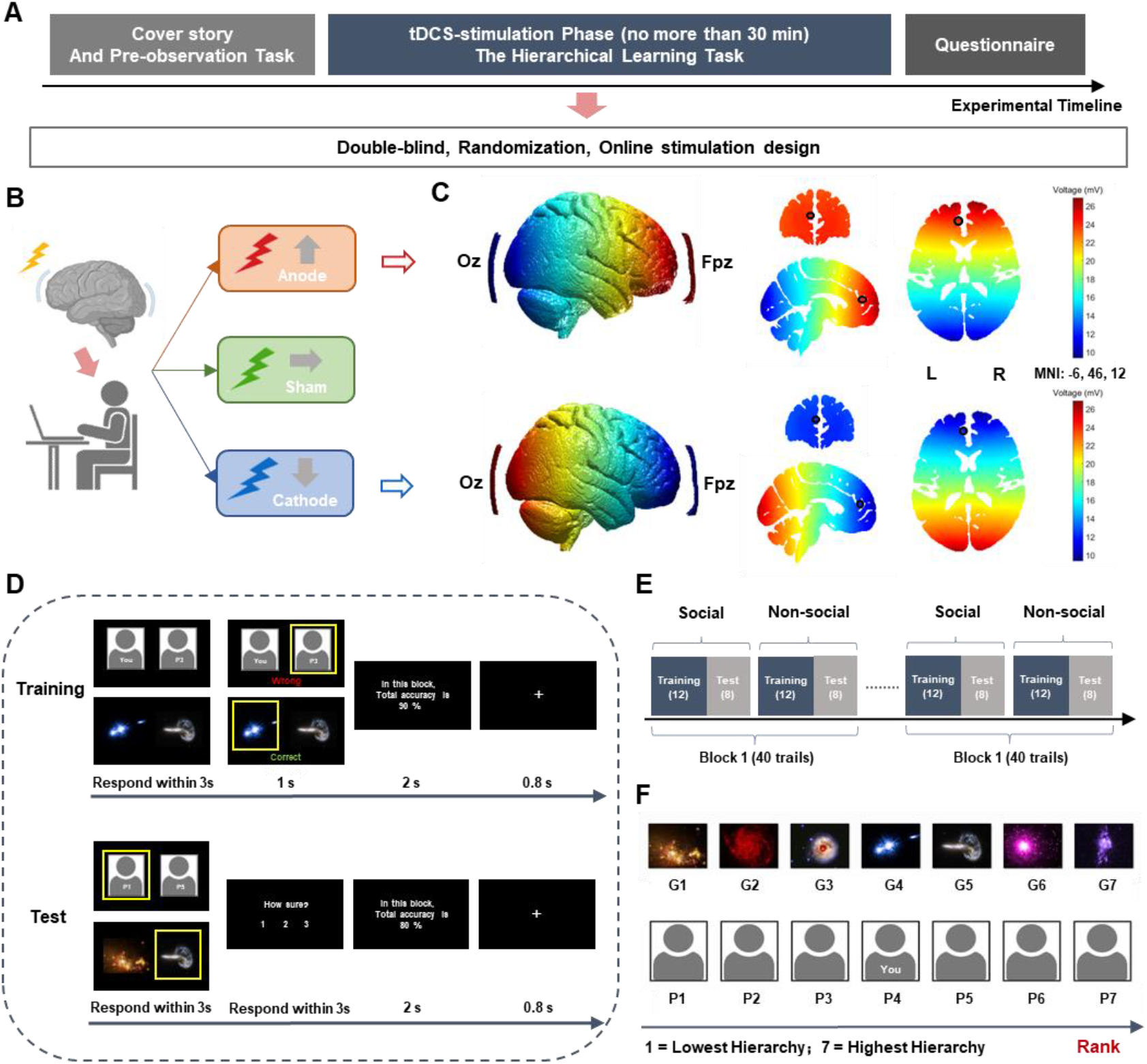
Illustration of the experimental procedure and behavioral paradigm. **A and B) Procedure.** First participants were given the cover story. They were asked to imagine that they recently had joined a technology company that detected precious minerals in different galaxies. As new members of the company, they were instructed to learn the hierarchical relationships between the staff (Social) and the mineral contents of the different galaxies (Non-social). To familiarize themselves with the company members and galaxies, they were instructed to passively observe all the pictures (7 faces, 7 galaxies) in an observation task. Each picture was randomly presented three times. Next, participants were randomly assigned to the Anode, Sham, or Cathode stimulation, and instructed to perform the task. At the end of the experiment, participants were required to complete questionnaires [see Method]. **C) Electric field simulation results for anodal and cathodal transcranial direct-current stimulation (tDCS).** The Fpz-Oz montage was chosen based on previous studies as targeting mPFC. The Simulated voltage distribution over the prefrontal cortex (left), and in coronal, sagittal, and axial slices (right) using the anodal montage with the MNI template brain. The black circle shows the targeted mPFC coordinates from Kumaran et al. (2016) (MNI: -6, 46, 12). The voltage indicates the strength of tDCS across the brain. L=left; R=right. **D and E) Hierarchy Learning Task.** There were 12 blocks including 12 training trials and 8 test trials. The Non-Social condition was identical to the Social condition except the stimuli were pictures of galaxies. **Training phase:** Participants were presented with adjacent items of the hierarchy (e.g., P4 vs P5, G4 vs G5, where P4 = “You”; and G4 = galaxy of rank equal to 4). They had to indicate the person they thought had higher status or the galaxy with more minerals. Through the correct feedback on their selection, they were able to learn the hierarchical relationships between the adjacent items. **Test phase:** Participants were required to view non-adjacent items in the hierarchy (e.g., P1 vs P5, G1 vs G5), infer which was the higher-ranked item, and rate the confidence of their selection—no feedback was provided. **F) Stimuli.** For Social condition, human faces were gender matched (here we used silhouette as illustration). For Non-social conditions, galaxy pictures were the same for females and males (1=Lowest position, 7=highest).

**Figure 2.**
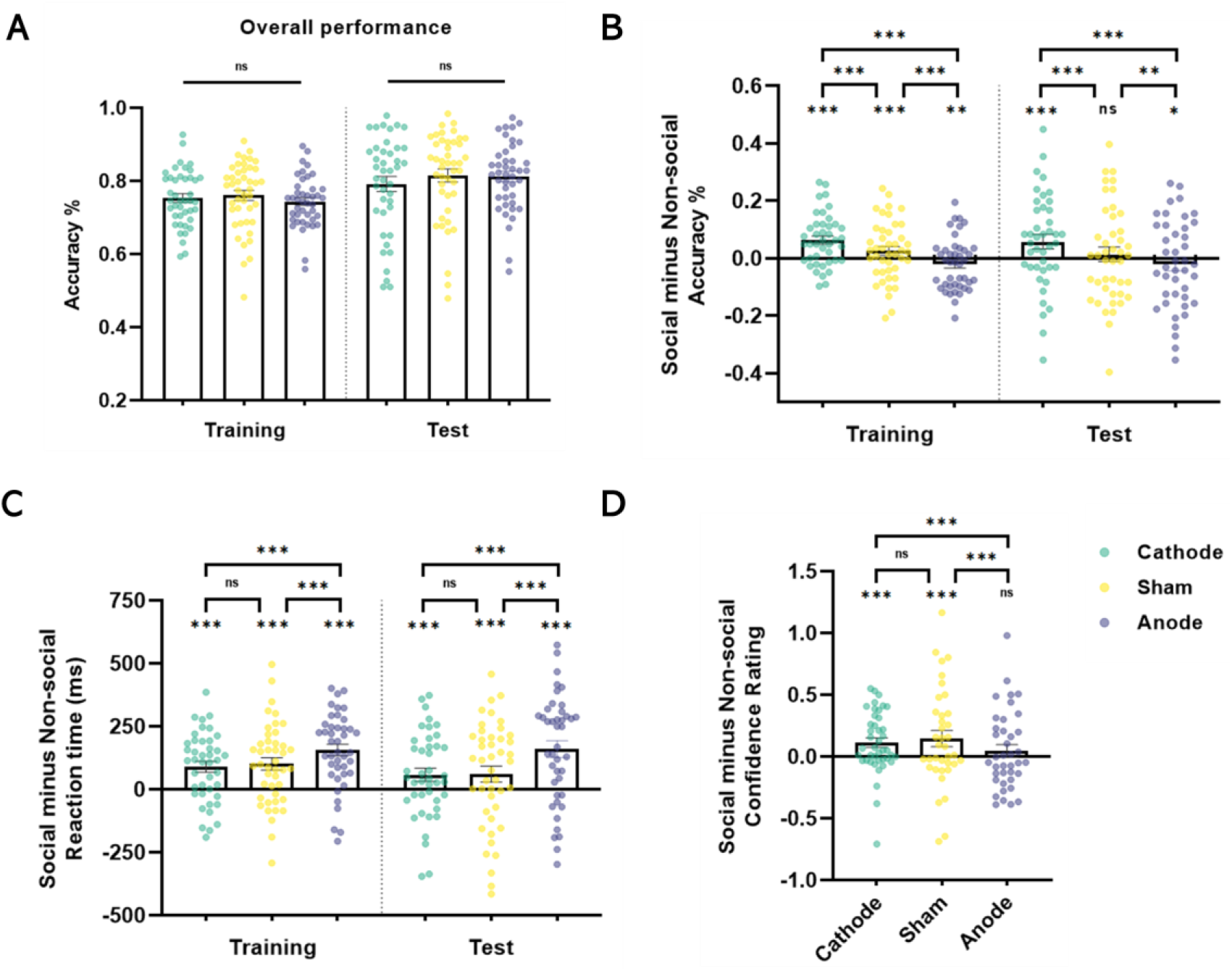
tDCS modulation in Training and Test phase. A) Overall performance combining both social and non-social condition; Social compared to Non-Social hierarchy learning B) performance accuracy %, C) reaction time (ms), and D) confidence rating of transitive inference decisions. The significance label on the top of the bars indicates the comparison between Social and Non-Social conditions. The significance label beneath the bars indicates the comparison between Cathode, Sham, and Anode stimulation of the differences between Social and Non-Social conditions. The significance levels were estimated by the marginal average effect across all individuals and time periods in the panel dataset. (*indicates *P*<0.05, **indicates *P*<0.005, ***indicates *P*<0.001; ****indicates *P*<0.0001; ns indicates non-significant; Error bars show SEM)

## Results

To investigate how brain stimulation modulated hierarchy learning, we conducted panel logistic or linear regressions, depending on the form of dependent variable (accuracy, reaction time, or confidence rating for each trial), on the population-average (generalized estimating equation, GEE) [see Methods for details of statistics]. These analyses allowed us to observe the effect of stimulation at the level of the population, by considering the effect of time. The independent variables were the tDCS stimulation (Anode/Sham/Cathode), hierarchy condition (Social/Non-social), and block number (1-12). The percentage change effect was estimated via the marginal effect (reported as *β* value below, see Methods). The marginal estimation calculates the average effect across all individuals and time periods in the panel dataset, which can easily be interpreted as the discreet change of the dependent variable given a unitary change of an independent variable.

### Effect of tDCS on Learning Hierarchical knowledge (Training Phase)

We first focused on the impact of mPFC-targeted brain stimulation regimes on hierarchical learning behavior between Social and Non-Social conditions. During the training phase, the overall learning accuracy was 0.752 ± 0.080 *SD* (Sham=0.760 ± 0.092 *SD*, Cathode=0.753 ± 0.076 *SD*, Anode=0.743 ± 0.069 *SD*). As shown in Fig.2A, there is no significant different in the overall performance when combining both social and non-social conditions. Panel logistic regression shows, under both Cathode and Sham stimulation, participants learned better in the Social condition relative to the Non-social [Probability of accuracy Cathode Social>Non-social: *β* = 0.061, SE = 0.007, z = 8.27, *P*<0.001, 95% CI (0.046, 0.075); Sham Social>Non-social: *β* = 0.024, SE = 0.070, z = 3.52, *P*<0.001, 95% CI(0.011, 0.038); Fig.2B]. Moreover, cathodal stimulation increased social hierarchy learning compared to sham stimulation [Contrasts of average marginal effects Cathode>Sham in Training: *χ*^2^ (1) = 12.68, *P* <0.001]. On the contrary, anodal stimulation significantly decreased accuracy in the Social condition compared to Non-Social [Anode Social<Non-social: *β* = -0.021, SE = 0.007, z = -2.80, *P* =0.005, 95% CI(-0.035, -0.006); Contrasts of average marginal effects Anode<Sham in Training: *χ*^2^ (1) = 19.82, *P*<0.0001; Fig.2B]. Furthermore, participants spent significantly more time to make decisions when facing the social hierarchy compared to non-social under anodal stimulation [Reaction Time Anode>Sham in Training: *χ*^2^(1)=25.44, *P*<0.0001; Fig.2C], suggesting an impairment of updating/learning social hierarchy knowledge under anodal stimulation.

### Effect of tDCS on Making Hierarchical Transitive Inference (Test Phase)

During the test phase, the overall transitive inference accuracy was 0.806 ± 0.116 *SD* (Sham=0.815 ± 0.119 *SD*, Cathode=0.792 ± 0.132 *SD*, Anode=0.812 ± 0.095 *SD*). There was no significant different in the overall performance when combining both social and non-social conditions (see Fig.2A). As shown in Fig.2B, under anodal stimulation participants’ performance to infer social hierarchy was significantly worse than on the non-social hierarchy task [Anode Social<Non-social: *β* = -0.020, SE= 0.008, z= -2.53, *P*<0.05, 95% CI(-0.036, -0.005); Contrasts of average marginal effects Anode<Sham in Test: *χ*^2^(1)=9.25, *P*<0.005], in contrast, performance to infer social hierarchy was significantly better under cathodal stimulation [Cathode Social>Non-social: *β* = 0.055, SE = 0.008, z= 6.49, *P*<0.001, 95% CI(0.038, 0.072); Contrasts of average marginal effects Cathode>Sham in Test: *χ*^2^(1)=13.39, *P*<0.0005].

However, under sham stimulation, unlike during training trials, there was no difference in performance accuracy between social and non-social hierarchy learning when making transitive inference [Sham: *β* = 0.013, SE = 0.008, z = 1.76, *P* =0.078, 95% CI(-0.001, 0.028)]. Similar to the Training phase, anodal stimulation significantly slowed the making of transitive inferences in Social compared to Non-social hierarchy [Reaction time Anode Social>Non-social: *β* = 159.783, SE = 9.861, z = 16.20, *P* <0.0001, 95% CI(140.456, 179.109); Contrasts of average marginal effects Anode>Sham in Test: *χ*^2^ (1) = 52.2, *P*<0.0001; Fig.2C].

During Test trials, participants were also required to rate the confidence in their choices. We expect participants to show higher confidence rating in social hierarchy inference compared to the non-social condition. The analysis of confidence ratings in the transitive judgements showed that under cathode and sham stimulation, in line with our hypothesis, participants were more confident in their Social than Non-Social hierarchy decisions [Cathode Social>Non-social: *β*=0.129, SE = 0.012, z = 10.6, *P* <0.001, 95% CI(0.105, 0.153); Sham Social>Non-social: *β* = 0.155, SE = 0.013, z = 11.94, *P* <0.001, 95% CI(0.130, 0.181); Fig.2D]. This was not the case for the Anodal stimulation group [Anode Social > Non-social: *β* = 0.022, SE = 0.127, z = 1.74, *P* =0.081, 95% CI(-0.003, 0.047)]. Moreover, in the Social compared to Non-Social conditions, participants under anodal stimulation were less confident in their judgments compared to Sham [Contrasts of average marginal effects Anode<Sham: *χ*^2^ (1) = 53.86, *P* <0.0001; Fig.2D]. These results are consistent with the worse performance observed under anodal tDCS in the social condition compared to the non-social, suggesting that participants were aware of their poorer performance in this condition.

Overall, the above findings indicate that the mPFC-targeted stimulation has different impacts during social and non-social hierarchy learning. Cathodal stimulation improved social hierarchy learning whereas anodal stimulation impaired it. Without tDCS (sham condition), participants learned social hierarchies better than non-social hierarchies, but there was no significant effect on transitive inferences.

### Anode Stimulation Impairs Hierarchy Learning in the Social Condition

We estimated the marginal effect of learning a Social or Non-Social hierarchy under each specific mPFC-targeted stimulation compared to Sham condition. This analysis was performed on the performance of Training and Test phases independently. In line with prior findings, compared to Sham, Anodal stimulation resulted in significantly lower accuracy during social hierarchy learning in both Training and Test trials [Social Training Anode<Sham: *β* = -0.045, SE = 0.018, z= -2.45, *P*=0.014, 95% CI (-0.08, -0.009); Social Test Anode<Sham: *β* = -0.052, SE=0.025, z= -2.07, *P*=0.038, 95% CI (-0.102, -0.003); Fig.3A]. Cathodal stimulation showed no significant effect on social hierarchy learning [Social Training Cathode<Sham: *β* =0.017, SE = 0.017, z = 0.99, *P* =0.322, 95% CI(-0.017, 0.05); Social Test Cathode<Sham: *β* = 0.002, SE = 0.025, z = 0.07, *P* =0.941, 95% CI(-0.047, 0.05)], and there was no significant impact of either anodal or cathodal tDCS during the learning of non-social hierarchies [Fig.3A]. Moreover, brain stimulation showed a selective effect on the reaction time during the Test but not the Training phase, and only in the Social condition [Social Test Anode>Sham: *β* = 90.619, SE= 38.061, z= 2.38, *P*=0.017, 95% CI(16.021, 165.2161); Fig.3B]. Combined with the findings on learning performance, these results substantiate that the mPFC-targeted tDCS stimulation modulates social hierarchy learning, but not non-social hierarchy and that anodal stimulation impairs the performance of learning social hierarchy.

**Figure 3.**
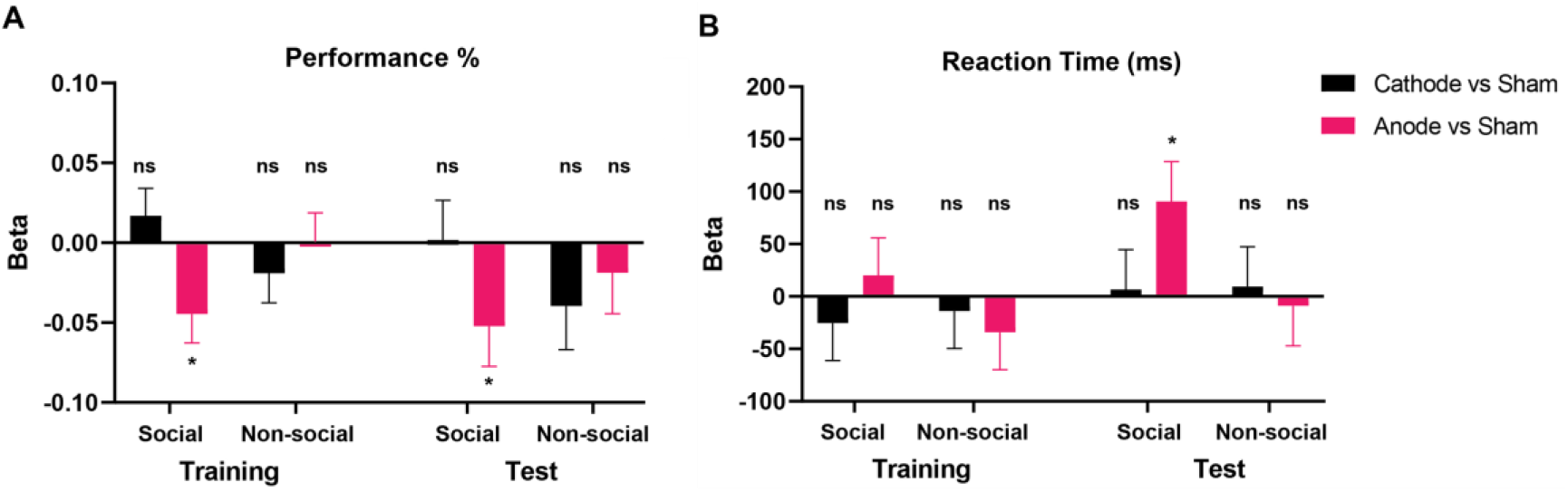
Brain stimulation selectively modulates social hierarchy learning during Training and Test phase A) performant accuracy % and B) reaction time (ms). The significance label on top or underneath the bars indicates the comparison between Cathode and Sham, or Anode and Sham. The y-axis indicates the estimated marginal effect of the linear model. The significance levels were estimated by the average marginal effect across all individuals and time periods in the panel dataset. (*indicates *P*<0.05, **indicates *P*<0.005, ***indicates *P*<0.001; ****indicates *P*<0.0001; ns indicates non-significant; Error bars show SEM)

### tDCS Impact on Block-to-Block Social Hierarchical Learning Performance

Next, we explored the effect of tDCS on the rate of learning hierarchical knowledge from trial block to trial block. We estimated the marginal effect of the block-to-block slope, in which the significant estimation reveals an increase or decrease of speed of learning (see Figure 4). There were contrasting effects of stimulation in the Training and Test phases.

**Figure 4.**
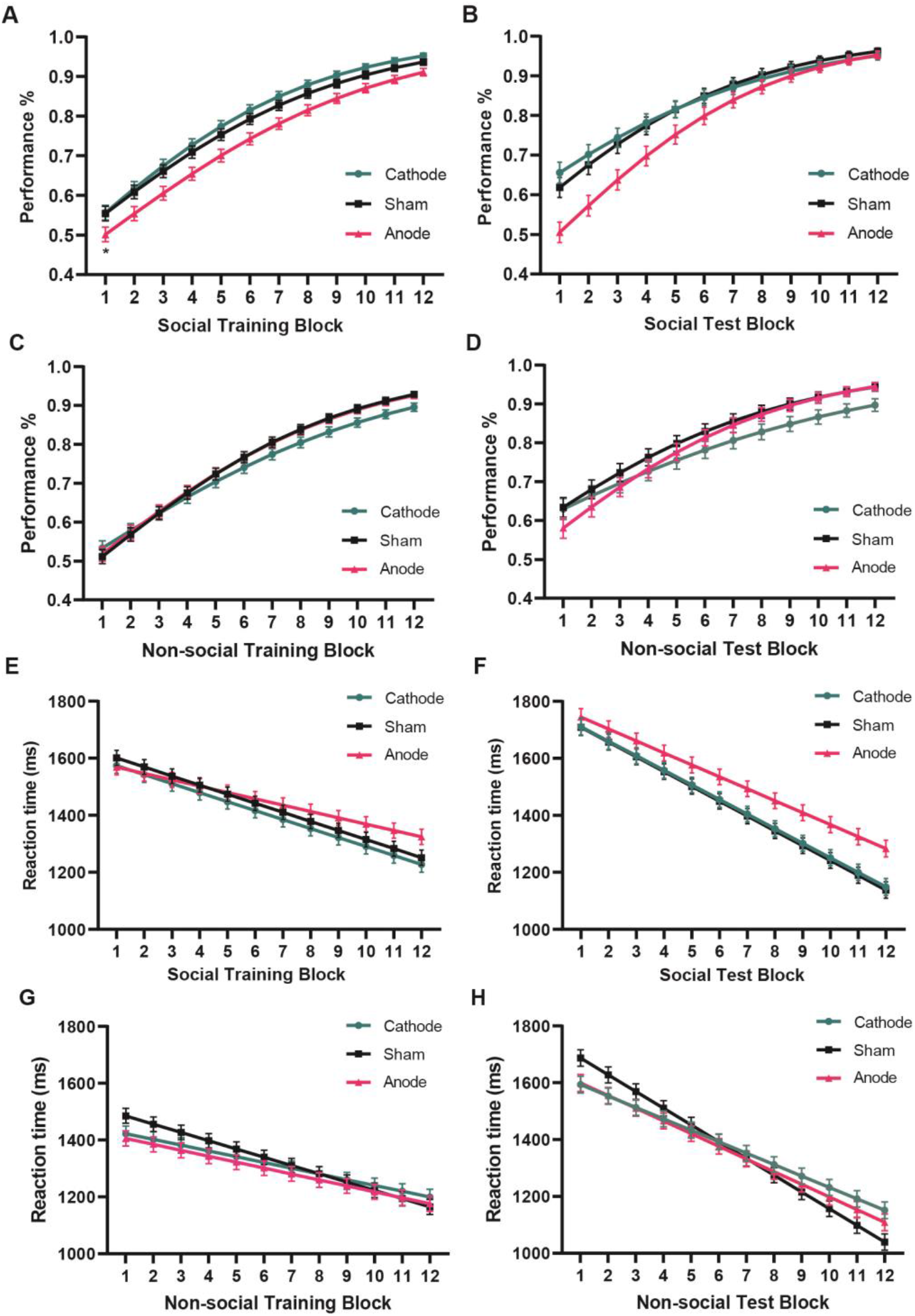
Anodal tDCS modulates social hierarchy learning from trial block to block. Modulation of the evolution of **the success rate, from trial block to trial block** by tDCS over consecutive **Social Training trial blocks (A)**, **Social Test trial blocks (B), Non-social Training trial blocks (C)**, **Non-social Test trial blocks (D)**. Modulation of the evolution of Reaction Time reduction from trial block to trial block by anodal tDCS. Anodal tDCS results in a reduction in the rate at which reaction time is reduced over consecutive trial blocks in both the **Social Training (E) and Test trial blocks (F**), as well as **Non-social Training (G) and Test trial blocks (H).** See Table S3 -S6 for the detailed significance slope comparison results of each block. (Error bars show SEM)

First, there was a significant improvement in performance from block to block over consecutive blocks in both phases [Training block: *β* = 0.035, SE = 0.001, z = 54.88, *P*<0.0001, 95% CI(0.035, 0.037); Test block: *β* =0.031, SE =0.001, z = 35.78, *P*<0.0001, 95% CI(0.029, 0.032)]. This effect confirms that participants were learning the hierarchies and were efficiently building on this learning to make successful transitive inferences as they progressed in the blocks. Note that the overall performance in the training phase is 0.752±0.080 SD regardless of condition, while the performance in the test phase is 0.806 ±0.116 SD. This difference between training and test performance was significant (P<0.001), perhaps due to the design of the test trials. Indeed, for 6/7 of the elements, they either always win in every trial in which they are involved (i.e., P1, P2 and P3) or always lose ( P5, P6, P7). The only element that sometimes wins and sometimes loses was P4 (or G4 for non-social condition). This may have facilitated performance on the test trials. To test this, we conducted an analysis of test trials by separating those trials with and without P4/G4. We found the overall accuracy was significant lower in the trials involved P4/G4 compared to the trials without P4/G4 (t=-2.233, P=0.027). This may explain why the performance of the test trials is higher than the performance of the learning trials which included a higher proportion of trials containing “ambiguous” items that were sometimes lower status and sometimes higher depending on the item with which they were paired.

Second, there was no significant effect of tDCS on overall hierarchical learning performance during the trial and error Training blocks [same slopes in accuracy rate (block) estimated function; Cathode vs Sham: *χ*^2^(1)=1.35, *P*=0.24, Anode vs Sham: *χ*^2^(1)=0.21, *P*=0.65]. However, there was a significant effect of anodal stimulation, which increased the rate of acquisition of the ability to make transitive inferences from the learned trial and error trials, when compared to sham tDCS, and cathode stimulation induced the opposite effect [Test block Slope Anode>Sham: *β* = 0.007, SE=0.002, *χ*^2^(1)=10.72, *P*=0.0008, 95% CI(0.003, 0.011); Slope Cathode>Sham: *β* = -0.004, SE=0.002, *χ*^2^(1)=3.70, *P*=0.046, 95% CI(-0.008, -0.00008)]. These results show that mPFC stimulation did not affect the speed of learning adjacent stimuli (Training phase), but rather the use of previously acquired knowledge required to make transitive inferences of hierarchy.

Furthermore, when we investigated social and non-social learning separately, we found that anodal stimulation specifically impacted the accuracy of the making of transitive inferences in the social condition by improving the accuracy of performance from block to block [Test block Slope Anode>Sham: *β*=0.009, SE=0.003, *χ*^2^(1)=11.82, *P*=0.0006, 95% CI(0.004, 0.014); Slope Cathode>Sham: *β*=-0.004, SE=0.003, *χ*^2^(1)=2.74, *P*=0.098, 95% CI(-0.010, 0.0008)], especially during earlier blocks. [Fig.4B and Table S3, for Non-social see Fig.4D and Table S5]. However, this effect appears to be the product of a generally worse performance during the training phase in each block, rather than any adverse effect on the making of transitive inferences per se. Indeed, the improved performance in making transitive inferences from block to block after anodal stimulation simply reflects that there is more room for improvement because of the lower performance during the preceding training trial and transitive inference blocks [Fig.4A and Table S3].

More intriguingly, in the Social condition, anodal stimulation also resulted in a decrease in the rate of reduction of reaction times from trial block to trial block during both the Training and Test phase, compared to Sham, whereas cathodal stimulation resulted in no significant change [Social Training: Slope anode>sham: *β*=9.580, SE=2.207, *χ*^2^(1)=18.84, *P*<0.0001, 95% CI(5.254, 13.906); Slope cathode>sham: *β*=0.285, SE=2.207, *χ*^2^(1)=0.02, *P*=0.897, 95% CI(-4.041, 4.611), Fig.4E; Social Test: Slope anode>sham: *β* = 9.970, SE = 2.824, *χ*^2^ (1) =12.47, *P*=0.0004, 95% CI(4.435, 15.505); Slope cathode>sham: *β* = 0.791, SE = 2.824, *χ*^2^ (1) =0.08, *P* =0.779, 95% CI(-4.743, 6.326), Fig.4F; see Table S4 for Social and Table S6 and Fig. 4G-H for Non-social]. This further suggests that anodal mPFC stimulation influences the modulation of social hierarchical knowledge updating and transitive inferences differently.

### mPFC-targeted Anode tDCS has Different Impacts on Learning Based on Social Ranks

Finally, to explore whether the causal role of mPFC on learning social hierarchy is influenced by relative social rank (i.e. higher or lower status). We thought to split trials according to those involving higher hierarchical status (Trials included Social: P4, P5, P6, P7; Non-social: G4, G5, G6, G7) and lower hierarchical status (Trials included Social: P1, P2, P3, P4; Non-social: G1, G2, G3, G4) in the Training phase. As shown in Fig.5A, the tDCS-induced deficit in social hierarchy learning impinged asymmetrically on trials involving higher social ranks [Social higher ranks Anode<Sham: *β* = -0.055, SE= 0.020, z = -2.69, *P* =0.007, 95% CI (-0.095, -0.015)], with no significant effect on trials involving lower social ranks [Social lower ranks Anode vs Sham: *β* = -0.030, SE = 0.021, z = -1.44, *P*=0.150, 95% CI(-0.070, 0.011)]. No similar asymmetrical effect was observed with respect to reaction times [Fig.5B]. This effect of anodal tCDS cannot be accounted for by self-involvement or related factors because it remained robust when we restricted our analysis to those trials that only involved the knowledge updating concerning other individuals in the hierarchy [i.e., trials that included the participants themselves were excluded (P4, and G4 in Non-social); Social training Accuracy Anode<Sham: *β* = -0.053, SE =0.020, z = -2.69, *P* =0.007, 95% CI(-0.091, -0.014); Social test Accuracy Anode<Sham: *β* = -0.047, SE =0.026, z = -1.83, *P* =0.068, 95% CI(-0.097, 0.003)]. These results imply that the mPFC updated social hierarchy information concerning members of the social hierarchy of higher status than oneself. Moreover, as shown in Table S2, there was no significant difference in age, choice bias, belief in cover story manipulation, sensation of tDCS stimulation, and social dominance orientation among the three tDCS stimulation groups. Thus, any effect of the groups on social hierarchy learning behavior cannot be accounted for by the preexisting group differences or sensation of the current stimulus.

**Figure 5.**
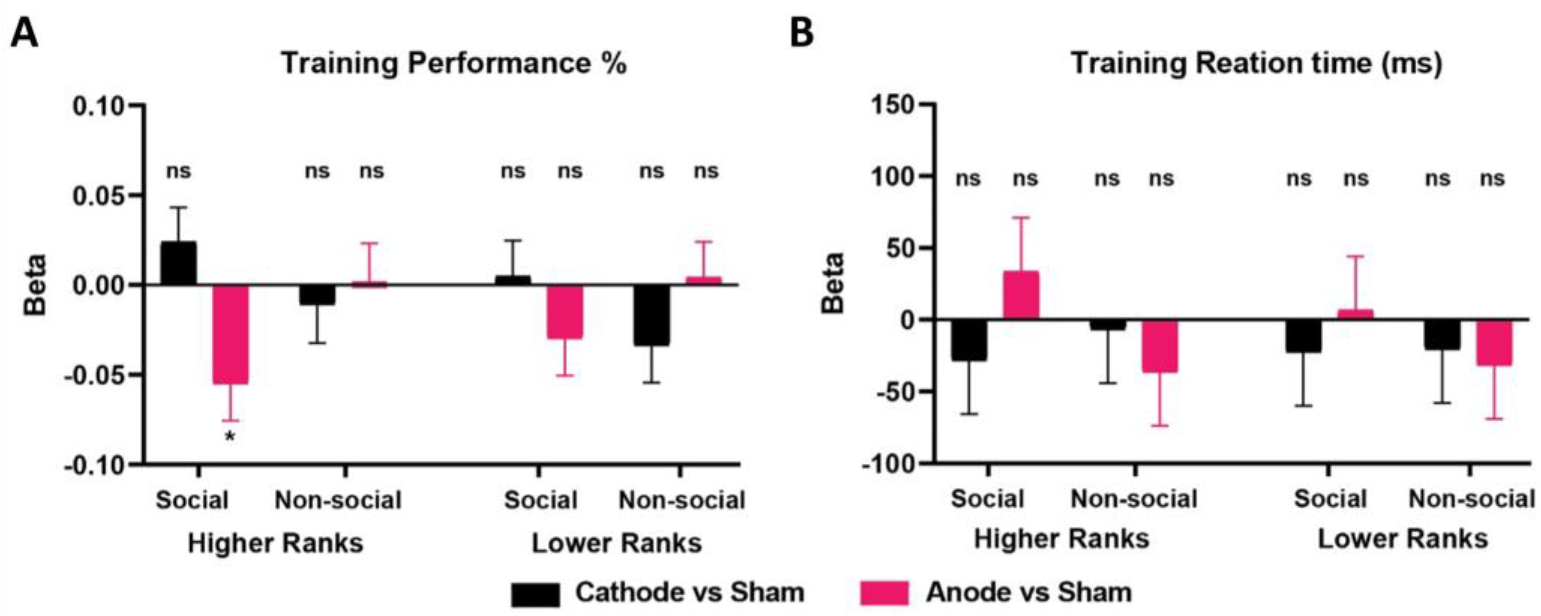
Anode tDCS selectively modulates social hierarchy learning A) performance accuracy % and B) reaction time (ms) based on social ranks. The significance label on the top of the bars indicates the comparison between Cathode and Sham, or Anode and Sham. The y-axis indicates the estimated marginal effect of the linear model. (*indicate *P*<0.05; **indicate *P*<0.01; ***indicate *P*<0.001; ns indicates non-significant; Error bars show SEM)

Taken together, these results suggest a causal role of mPFC in tracking the development of knowledge about Social, but not Non-social hierarchy. Anodal tDCS on the mPFC impaired social hierarchy learning performance. The anodal tDCS had distinct effects on the learning of social hierarchy knowledge and the making of transitive inference judgments that appear to be specific to hierarchy members of higher status.

## Discussion

Our study examined the causal role of the mPFC in learning and making transitive inferences about social and non-social hierarchy relationships using tDCS stimulation. Anodal stimulation over the mPFC modulated social but not non-social hierarchy learning, which provides causal evidence implicating the mPFC in the establishment of *social* hierarchy knowledge. The two-phase hierarchy learning task adopted in our study allowed us to effectively separate the updating and confirmation of hierarchical knowledge during the trial and error learning of the hierarchy in each Training block from the making of transitive inferences, on the basis of the acquired information, during the Test phase of each block. Anodal tDCS appeared to reduce the global performance of building and updating the social hierarchy knowledge during Training phase. Mechanically, because learning of the social ranks was not established during the early blocks of Training phase, this also reduced the success of transitive inference making during Test phases in the anodal group [Fig.2]. However, during later blocks of the test phase, when learning of the social hierarchy improved, the detrimental effect of anodal tDCS gradually decreased and disappeared [Fig.3B]. This shows that anodal tDCS does not disturb the making of inferences for social hierarchy when the learning of social ranks had improved over blocks. The apparent decrease in accuracy observed in the early blocks of the test phase in the Anodal group in social vs non-social hierarchies is probably only the consequence of the impaired learning of social hierarchy during early training [Fig.3A]. Thus, mPFC anodal stimulation disturbs the learning of social hierarchy, but not the making of transitive inferences from it.

Our findings that anodal mPFC stimulation disrupts the learning of social hierarchies, but leaves intact the learning of non-social hierarchies, indicate that the learning of these two types of hierarchies may rely on distinct cognitive processes. The fact that social hierarchy depends causally on the mPFC resonates with recent studies suggesting that task representations may differ across domains, such as the spatial and conceptual domains ^33^, or abstract vs naturalistic domains ^34^. Here, the nature of the items themselves (faces vs galaxies) may have influenced how they were learned because faces may be more easily learned. Confirming this hypothesis, differences in both learning accuracy and confidence were observed when directly comparing the sham group in the Social and Non-social conditions.

Another important finding from our study is that mPFC stimulation left the transitive inferences making processes unimpaired. This indicates that the mPFC is not causally necessary to make transitive inferences. A recent simulation of electrode fields has shown that our montage of Fpz-Oz generates higher electric currents in the amygdala and the hippocampus, which might facilitate the modulation of deep brain regions ^35^. The impact of the mPFC anodal stimulation on learning social hierarchy behavior may not be mediated by local activity alone, but by directed communication with other brain areas. For example, the hippocampus has been reported to encode abstract general knowledge of relationships whatever their nature (i.e. spatial, social, and non-social) ^36^. It is not only involved in forming the cognitive maps to organize information simultaneously ^37,38^, it also contributes to concept learning by representing the feature combinations related to current behaviors ^39,40^. The mPFC represents stimuli-outcome relationships of the cognitive map ^41^ and receives input information from hippocampus to update current information ^42^. Consistent with this, the mPFC was found to selectively mediate the learning of knowledge about social hierarchy whereas domain-general coding of ranks was observed in the hippocampus, even when the task did not require it ^15^. Although another fMRI report did not find mPFC in social vs non-social hierarchy learning^16^, this was only a correlational fMRI study, which cannot account for the causal role of the mPFC. Functional coupling between the mPFC and hippocampus has been shown to support social learning and is also involved in both conceptual learning and episodic memory of cognitive maps ^43,44^. This may explain why mPFC-targeted tDCS stimulation perturbed the social hierarchy knowledge updating across blocks, but only impaired the transitive inference during the earlier blocks when the conceptual knowledge was not yet well established.

Our results show that anodal tDCS perturbates performance in an asymmetrical manner and preferentially impinges on social comparison processes concerning those of status superior to one’s self, but having no significant effect on those ranked below. The social comparison theory posits that people are driven to compare themselves to others for accurate self-evaluations ^29^. Specifically, people compare themselves to others in two opposite directions -downward and upward- that differ in motivations, comparison targets, and consequences ^30,31^. The upward comparison refers to comparing those who are thought to rank higher. Upward comparison is most likely performed to fulfill the motivation of challenging others and self-improvement. This type of social comparison invokes threat to the self ^45^ and provokes negative emotions such as envy ^46,47^. In contrast, the downward comparison is most likely done to fulfill the motivation of self-enhancement. A previous study also suggest the causal role of dmPFC in self-other mergence^48^. Our findings that the mPFC is engaged or focused on individuals ranking higher than oneself demonstrate that this region is causally necessary for upward comparison. Our results agree with fMRI findings reporting that mPFC distinguishes between higher and lower ranks with respect to oneself ^15^, and engagement in different types of social valuation processes^49,50^.

Overall, our tDCS approach establishes a causal relationship between mPFC and social hierarchy learning. The maladaptive assessment of social dominance hierarchies is an important source of distress in social disorders such as depression and anxiety ^10,51^. Our findings not only extend our understanding of the role of mPFC and its involvement in social learning processes, they also suggest a possible novel avenue of treatment based on brain stimulation techniques to treat neuropsychiatric disorders (e.g., depression, anxiety) in which the experience of repeated social defeats often leads to social avoidance ^2,52^.

## Methods

### Participants

A total of 136 participants (67 males, 69 females) were recruited via online fliers with informed written consent. All participants were right-handed, with no history of psychiatric or neurologic disorders, and were randomly assigned to receive anode, cathode, or sham stimulation over the medial prefrontal cortex (mPFC) while performing the hierarchy learning tasks. We set a threshold of 80% accuracy in Training phase. Six participants did not reach this *priori* threshold and were excluded from the analysis because they did not learn either Social or Non-social conditions in the training trials sufficiently well (the accuracy rate of each block was lower than 2/3). In addition, one participant was excluded because he responded stereotypically (i.e., one key for the whole block), and another because the program was restarted twice. Thus, the data from 128 participants (males=63, mean age=19.90±0.145) were analyzed (Sham=44: Male=23, Female=21; Cathode=42: Male=21, Female=21; Anode=42: Male=21, Female=21). The study was approved by the ethics committee of South China Normal University and all participants received 45 CNY after the task.

### Stimuli

Images of faces in the Social condition were selected from the CAS-PEAL Large-Scale Chinese Face Database ^53^. Silhouettes (2 faces, 1 female, 1 male), were used to represent “You” which refers to the participant in the experiment. Frontal images (12 neutral faces, 6 females, 6 males) identified fictive hierarchy members for the subsequent experiments. The hair and neck were preserved for the facial pictures. For female participants, the hierarchy was composed of pictures of females and *vice versa* for male participants. Previous studies have shown facial gender can influence perceptions, judgments, and behavior related to social hierarchy and social dominance ^54–56^. By matching the gender between participants and stimuli we sought to ensure that participants perceived and evaluated the facial stimuli consistent with their own gender-related expectations, to reduce the number of potential factors that might influence the effect of social hierarchy learning and tDCS stimulation. Images of galaxies were selected from a public astronomy website (http://hubblesite.org/gallery/album/nbula). All pictures were processed by Adobe Photoshop software to ensure grayscale and resolution were consistent. The hierarchical ranks of galaxies and face stimuli were randomized across the groups. The experiment was programmed in E-prime 2.0 and presented on a 14-inch laptop.

### Experiment Procedure

With a double-blind and sham-control design, our study included three phases: Cover story and Pre-observation Task, tDCS stimulation phase (Hierarchical Learning Task), and Questionnaires [Fig.1].

### Cover story

Participants were asked to imagine they had joined recently a technology company that detected precious minerals in different galaxies. Then, they were instructed to observe the photos of staff members and galaxies related to the company to familiarize themselves with company members and business (see Pre-observation task below). After the observation task, they were informed that as a new member of the company, they needed to learn the rank relationship between staff members to help them adjust to work.

In the meantime, they also needed to learn the relative mineral contents of different galaxies. The learning task included two phases. During a training phase, within each trial, they will see a pair of pictures (i.e. faces or galaxies), they will need to choose which one has more power or mineral, and following the decision they will have feedback of their choice. During a test phase, within each trial, they will again see a pair of pictures, however, there will be no feedback of their choice provided and they need to rate their confidence in the choice from 1 to 3. At the end of each block, they have feedback on the accuracy rate of the current block. They were informed that the paired items in the training and test phase are different, their task is to learn as much good as they can in both training and test phases.

### Pre-observation task

The observation task was to reduce the differential effects of extraneous stimuli during subsequent tasks. Participants were instructed to passively observe the pictures presented on the screen to familiarize themselves with the staff members in the company and the galaxies related to the company business. There was a silhouette that represents “You” which refers to the participant in the experiment. There were three blocks of the task. Each block included 14 trials (7 face pictures, 7 galaxy pictures; randomly presented). Within each trial, following an 800ms attention cross, the picture was presented for 3 seconds on the screen. Thus, participants observed each picture three times.

### Hierarchy Learning Task (performed under sham or tDCS stimulation)

During tDCS stimulation, participants were required to perform a Hierarchy Learning task, including Training and Test phases in both Social and Non-Social conditions. The condition presented first was consistent with the observation task and balanced pseudo-randomly among the participants. The sequences of paired pictures were randomized, as was the left or right location in which pictures were presented. Human faces (sex matched) were used in the Social condition, whereas images of galaxies were used in Non-Social condition In the Training Phase, participants were required to view a pair of adjacent hierarchical pictures [P4 vs P5, G4 vs G5; P=person and G=galaxy; P4 means “YOU”; Fig.1A] and identify which picture they thought had a higher rank (Social) or more minerals (Non-Social). They received accurate feedback of the correctness of their selection, and were thus able to learn the hierarchical relationships between the adjacent items. There were 12 blocks of Training phase, with each block including 2 six trial mini-blocks composed of the 6 paired items (P1 vs P2, P2 vs P3, P3 vs P4, P4 vs P5, P5 vs P6, P6 vs P7). Each Training trial block was followed by a Test trial block.

For the Test Phase blocks, hierarchically non-adjacent pairs of pictures were presented [e.g., P1 vs P5, G1 vs G5; Fig.1A]. Participants were required to make transitive inference judgments and rate the confidence of their decisions from 1 (guess) to 3 (very sure). There were 12 blocks of Test phase and each included a single 8-trial mini-block composed of 8 paired items [P1 vs P4, P2 vs P4, P2 vs P5, P2 vs P6, P3 vs P5, P3 vs P6, P4 vs P6, P1 vs P7]. For both Training and Test trials, at the end of each block, they received average accuracy feedback on their decisions.

### Questionnaires

After the Hierarchy learning task, participants were required to fill in the Social Dominance Orientation (SDO) scale, and answer the post-questions about the task and tDCS stimulation rating: i) the discomfort of electrode stimulation (from 1 for none to 5 for very discomforting); ii) how much they believed that they were one of the members of the company (from 1 for none to 10 for complete belief). A previous study found that the sensitivity of the right DLPFC to social ranks correlated positively with the SDO scale^57^, which is known to predict behaviors and political attitudes associated with the legitimization of dominance hierarchies^58^. Thus, the purpose of SDO measure was to control for any group difference in sensitivity to social rank.

### Brain stimulation and current modeling

We used NeuroConn transcranial direct current stimulation devices (NeuroConn, Ilmenau, Germany) for tDCS stimulation. According to previous studies which investigated the causal role of mPFC in social behaviors ^59,60^, we adopted the Fpz-Oz montage (EEG 10-20 system) with 1.5mA current as the stimulation protocol. The Fpz-Oz montage has been shown in a recent simulation of electrode fields study that not only targeted the mPFC regions but also generates higher electric currents in the amygdala and the hippocampus, which might facilitate the modulation of deep brain regions^35^. We used gel to improve conductivity and reduce skin irritation. The electrode sizes were both 5 cm x 7 cm (35 cm^2^) [Fig.1C]. In all stimulation conditions, the current intensity was 1.5 mA, applied with a 30-second fade-in and fade-out at the beginning, and the end of the stimulation. For both anode and cathode stimulation, the 1.5 mA stimulus lasted no more than 30 minutes (when participants completed the task in less than 30 minutes, the current was terminated earlier). For sham stimulation, the current only lasted 15 seconds. To account for possible delays in the onset of tDCS effects, participants were required to wait 2.5 minutes after the onset of stimulation to start the hierarchy learning task (thus, for the sham condition, the current stimulation had ceased since 2.25 minutes before the first learning trial).

To ensure that our electrode montage effectively stimulated the mPFC, electrical potential simulations were performed using ROAST ^61,62^ with the MNI152 template brain. Electrodes were simulated as pads, with a 70x50x3mm pad located over Fpz and Cz of standard 10-10 system locations. Tissue conductivities were set as white matter=0.11 S/m, gray matter=0.21 S/m, CSF=0.53 S/m, bone=0.02 S/m, and skin=0.90 S/m. For the anodal stimulation, 1.5mA was set as inward flowing current from the Fpz, and -1.5mA outward flowing current from the Cz. For the cathodal stimulation this was reversed.

### Statistical analyses

Behavioral analyses were conducted in STATA 14. For the panel regressions using the population-average effect (generalized estimating equation approach, GEE), this allowed us to estimate the effect of brain stimulation at the level of population and take into account the effect of time (Learning). The Panel data of t=480 trials clustered on each of i=128 participants were used.

We reported significant marginal effect of estimated β values. The model equation of GEE for panel logistic regressions:

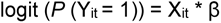

Yit represents the binary dependent variable for individual i at time t. Xit represents the vector of explanatory variables for individual i at time t. *P* represents the probability of a given event. β is the vector of coefficients to be estimated, representing the population-average marginal effects.

The marginal effect of the variable *X*it is estimated as:

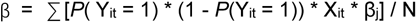

β represents the estimated marginal effect. *P*(Yit = 1) represents the predicted probability of the dependent variable being equal to 1 for individual i at time t. Xit represents the vector of explanatory variables for individual i at time t. βj represents the estimated coefficients from the GEE model. N represents the total number of observations in the panel dataset.

The marginal effect estimation equation calculates the average marginal effect across all individuals and time periods in the panel dataset. Thus, the estimated marginal effect can easily be interpreted as the discreet change of the dependent variable given a unitary change of an independent variable. It quantifies the change in the log-odds of the dependent variable associated with a one-unit change in the corresponding independent variable, while holding all other variables constant. In other words, it quantifies the impact of a unit change in the independent variable on the dependent variable, taking into account the effects of other variables in the model.

For the panel linear regressions, reported β value represents Xit, i.e. the regression coefficient. Indeed, in a linear regression, marginal effect of a variable is equal to the estimated coefficient.

## Data Availability

The behavioral data that support the findings of this study are available as a repository on the Open Science Framework: https://osf.io/zvcfh/

## Code availability

Software packages used for the analyses are STATA 14. Code used to generate the analyses are available on the Open Science Framework: https://osf.io/zvcfh/

## Acknowledgments

This research has benefited from the financial support of IDEXLYON from Université de Lyon (project INDEPTH) within the Programme Investissements d’Avenir (ANR-16-IDEX-0005) and of the LABEX CORTEX (ANR-11-LABX-0042) of Université de Lyon, within the program Investissements d’Avenir (ANR- 11-IDEX-007) operated by the French National Research Agency. This work was also supported by a grant (ADOSTRESS, ANR-21-CE37-0032-01) from the Agence Nationale pour la Recherche and a grant from the Fondation de France (FDF 00089590) to JCD, and a grant from National Natural Science Foundation of China to QC (31970982; 32171019). We thank Siying Li, and Yaner Su for helpful assistance with data collection.

## Author Contributions

C.Q. and J.-C.D designed research; C.Q., Y.H., S.C., and M.S. performed research; C.Q., Y.H., R.P., S.C., and J.-C.D analyzed data; E.D. and F.M. reviewed the paper; and C.Q., Y.H., R.P., and J.-C.D wrote the paper. All authors approved the manuscript final version.

## Additional information

Supplementary Information Text, Figure S1, and Tables S1 to S6

## References

1. Koski, J. E., Xie, H., & Olson, I. R. Understanding social hierarchies: The neural and psychological foundations of status perception. Social neuroscience, 10(5), 527–550 (2015).

2. Qu, C., Ligneul, R., Van der Henst, J. B., & Dreher, J. C. An integrative interdisciplinary perspective on social dominance hierarchies. Trends in cognitive sciences, 21(11), 893–908 (2017).

3. Munuera, J., Rigotti, M. & Salzman, C. D. J. N. n. Shared neural coding for social hierarchy and reward value in primate amygdala. 21, 415–423 (2018).

4. Ferreira-Fernandes, E. & Peça, J. J. F. i. C. N. The neural circuit architecture of social hierarchy in rodents and primates. 16, 874310 (2022).

5. Buston, P. Social hierarchies: Size and growth modification in clownfish. Nature 424, 145–146, (2003).

6. Grosenick, L., Clement, T. S. & Fernald, R. D. Fish can infer social rank by observation alone. Nature 445, 429–432, (2007).

7. Chiao, J. Y., Harada, T., Oby, E. R., Li, Z., Parrish, T., & Bridge, D. J. Neural representations of social status hierarchy in human inferior parietal cortex. Neuropsychologia, 47(2), 354–363 (2009).

8. Wang, F., Zhu, J., Zhu, H., Zhang, Q., Lin, Z., & Hu, H. Bidirectional control of social hierarchy by synaptic efficacy in medial prefrontal cortex. Science, 334(6056), 693–697 (2011).

9. Cheney, D. L., & Seyfarth, R. M. How monkeys see the world: Inside the mind of another species. University of Chicago Press (1990).

10. Sapolsky, R. M. The influence of social hierarchy on primate health. science, 308(5722), 648–652 (2005).

11. Cummins, D. D. How the social environment shaped the evolution of mind. Synthese, 122(1), 3–28 (2000).

12. Santamaría-García, H., Pannunzi, M., Ayneto, A., Deco, G., & Sebastián-Gallés, N. ‘If you are good, I get better’: the role of social hierarchy in perceptual decision-making. Social cognitive and affective neuroscience, 9(10), 1489–1497 (2014).

13. Feng, C., Li, Z., Feng, X., Wang, L., Tian, T., & Luo, Y. J. Social hierarchy modulates neural responses of empathy for pain. Social Cognitive and Affective Neuroscience, 11(3), 485–495 (2016).

14. Boyce, W. T. Social stratification, health, and violence in the very young. Annals of the New York Academy of Sciences, 1036(1), 47–68 (2004).

15. Kumaran, D., Banino, A., Blundell, C., Hassabis, D., & Dayan, P. Computations underlying social hierarchy learning: distinct neural mechanisms for updating and representing self-relevant information. Neuron, 92(5), 1135–1147 (2016).

16. Kumaran, D., Melo, H. L., & Duzel, E. The emergence and representation of knowledge about social and nonsocial hierarchies. Neuron, 76(3), 653–666 (2012).

17. Ligneul, R., Obeso, I., Ruff, C. C., & Dreher, J. C. Dynamical representation of dominance relationships in the human rostromedial prefrontal cortex. Current Biology, 26(23), 3107–3115 (2016).

18. Marsh, A. A., Blair, K. S., Jones, M. M., Soliman, N., & Blair, R. J. R. Dominance and submission: the ventrolateral prefrontal cortex and responses to status cues. Journal of cognitive neuroscience, 21(4), 713–724 (2009).

19. Todorov, A., Said, C. P., Engell, A. D., & Oosterhof, N. N. Understanding evaluation of faces on social dimensions. Trends in cognitive sciences, 12(12), 455–460 (2008).

20. Zink, C. F., Tong, Y., Chen, Q., Bassett, D. S., Stein, J. L., & Meyer-Lindenberg, A. Know your place: neural processing of social hierarchy in humans. Neuron, 58(2), 273–283 (2008).

21. Apps, M. A., & Sallet, J. Social learning in the medial prefrontal cortex. Trends in Cognitive Sciences, 21(3), 151–152 (2017).

22. Behrens, T. E., Muller, T. H., Whittington, J. C., Mark, S., Baram, A. B., Stachenfeld, K. L., & Kurth-Nelson, Z. What is a cognitive map? Organizing knowledge for flexible behavior. Neuron, 100(2), 490–509 (2018).

23. Gershman, S. J., & Niv, Y. Learning latent structure: carving nature at its joints. Current opinion in neurobiology, 20(2), 251–256 (2010).

24. Niv, Y. Learning task-state representations. Nature neuroscience, 22(10), 1544–1553 (2019).

25. Stachenfeld, K. L., Botvinick, M. M., & Gershman, S. J. The hippocampus as a predictive map. Nature neuroscience, 20(11), 1643–1653 (2017).

26. Whittington, J. C., Muller, T. H., Mark, S., Chen, G., Barry, C., Burgess, N., & Behrens, T. E. The Tolman-Eichenbaum machine: unifying space and relational memory through generalization in the hippocampal formation. Cell, 183(5), 1249–1263 (2020).

27. Baram, A. B., Muller, T. H., Nili, H., Garvert, M. M., & Behrens, T. E. J. Entorhinal and ventromedial prefrontal cortices abstract and generalize the structure of reinforcement learning problems. Neuron, 109(4), 713–723 (2021).

28. Park, S. A., Miller, D. S., Nili, H., Ranganath, C., & Boorman, E. D. Map making: constructing, combining, and inferring on abstract cognitive maps. Neuron, 107(6), 1226–1238 (2020).

29. Festinger, L. A theory of social comparison processes. Human relations, 7(2), 117–140 (1954).

30. Latané, B. Studies in social comparison—Introduction and overview. Journal of Experimental Social Psychology, 1, 1–5 (1966).

31. Wills, T. A. Downward comparison principles in social psychology. Psychological bulletin, 90(2), 245 (1981).

32. Brunoni, A. R. et al. A systematic review on reporting and assessment of adverse effects associated with transcranial direct current stimulation. Int J Neuropsychoph 14, 1133–1145, (2011).

33. Wu, C. M., Schulz, E., Garvert, M. M., Meder, B., & Schuck, N. W. Similarities and differences in spatial and non-spatial cognitive maps. PLoS computational biology, 16(9), e1008149 (2020).

34. Farashahi, S., Xu, J., Wu, S. W., & Soltani, A. Learning arbitrary stimulus-reward associations for naturalistic stimuli involves transition from learning about features to learning about objects. Cognition, 205, 104425 (2020).

35. Gomez-Tames, J., Asai, A., & Hirata, A. Significant group-level hotspots found in deep brain regions during transcranial direct current stimulation (tDCS): A computational analysis of electric fields. Clinical Neurophysiology, 131(3), 755–765 (2020).

36. Morton, N. W., & Preston, A. R. Concept formation as a computational cognitive process. Current Opinion in Behavioral Sciences, 38, 83–89 (2021).

37. Park, S. A., Miller, D. S., Nili, H., Ranganath, C., & Boorman, E. D. Map making: constructing, combining, and inferring on abstract cognitive maps. Neuron, 107(6), 1226–1238 (2020).

38. Theves, S., Fernández, G., & Doeller, C. F. The hippocampus maps concept space, not feature space. Journal of Neuroscience, 40(38), 7318–7325 (2020).

39. Davis, T., Xue, G., Love, B. C., Preston, A. R., & Poldrack, R. A. Global neural pattern similarity as a common basis for categorization and recognition memory. Journal of Neuroscience, 34(22), 7472–7484 (2014).

40. Mack, M. L., Love, B. C., & Preston, A. R. Dynamic updating of hippocampal object representations reflects new conceptual knowledge. Proceedings of the National Academy of Sciences, 113(46), 13203–13208 (2016).

41. Mack, M. L., Preston, A. R., & Love, B. C. Ventromedial prefrontal cortex compression during concept learning. Nature communications, 11(1), 1–11 (2020).

42. Wikenheiser, A. M., & Schoenbaum, G. Over the river, through the woods: cognitive maps in the hippocampus and orbitofrontal cortex. Nature Reviews Neuroscience, 17(8), 513–523 (2016).

43. Schlichting, M. L., Mumford, J. A., & Preston, A. R. Learning-related representational changes reveal dissociable integration and separation signatures in the hippocampus and prefrontal cortex. Nature communications, 6(1), 1–10 (2015).

44. Schlichting, M. L., & Preston, A. R. Hippocampal–medial prefrontal circuit supports memory updating during learning and post-encoding rest. Neurobiology of learning and memory, 134, 91–106 (2016).

45. Brickman, P., & Bulman, R. J. Pleasure and pain in social comparison. Social comparison processes: Theoretical and empirical perspectives, 149, 186 (1977).

46. Chester, D. S., Powell, C. A., Smith, R. H., Joseph, J. E., Kedia, G., Combs, D. J., & DeWall, C. N. Justice for the average Joe: The role of envy and the mentalizing network in the deservingness of others’ misfortunes. Social Neuroscience, 8(6), 640–649 (2013).

47. Jankowski, K. F., & Takahashi, H. Cognitive neuroscience of social emotions and implications for psychopathology: examining embarrassment, guilt, envy, and schadenfreude. Psychiatry and clinical neurosciences, 68(5), 319–336 (2014).

48. Wittmann, M. K., Trudel, N., Trier, H. A., Klein-Flügge, M. C., Sel, A., Verhagen, L., & Rushworth, M. F. (2021). Causal manipulation of self-other mergence in the dorsomedial prefrontal cortex. Neuron, 109(14), 2353–2361.

49. Kim, H. Stability or Plasticity?–A Hierarchical Allostatic Regulation Model of Medial Prefrontal Cortex Function for Social Valuation. Frontiers in neuroscience, 14, 281 (2020).

50. Lebreton, M., Jorge, S., Michel, V., Thirion, B., & Pessiglione, M. An automatic valuation system in the human brain: evidence from functional neuroimaging. Neuron, 64(3), 431–439 (2009).

51. Johnson, S. L., Leedom, L. J., & Muhtadie, L. The dominance behavioral system and psychopathology: evidence from self-report, observational, and biological studies. Psychological bulletin, 138(4), 692 (2012).

52. Sellaro, R., Nitsche, M. A., & Colzato, L. S. The stimulated social brain: effects of transcranial direct current stimulation on social cognition. Annals of the New York Academy of Sciences, 1369(1), 218–239 (2016).

53. Gao, W., Cao, B., Shan, S., Chen, X., Zhou, D., Zhang, X., & Zhao, D. The CAS-PEAL large-scale Chinese face database and baseline evaluations. *IEEE Transactions on Systems*, Man, and Cybernetics-Part A: Systems and Humans, 38(1), 149–161 (2007).

54. Bruce, V. & Young, A. W. Face perception. (Psychology Press, 2012).

55. DeBruine, L. M., Jones, B. C., Smith, F. G., & Little, A. C. (2010). Are attractive men’s faces masculine or feminine? The importance of controlling confounds in face stimuli. Journal of Experimental Psychology: Human Perception and Performance, 36(3), 751.

56. Rule, N. O., & Ambady, N. (2009). She’s got the look: Inferences from female chief executive officers’ faces predict their success. Sex Roles, 61, 644–652.

57. Ligneul, R., Girard, R., & Dreher, J. C. (2017). Social brains and divides: the interplay between social dominance orientation and the neural sensitivity to hierarchical ranks. Scientific Reports, 7(1), 45920.

58. Pratto, F., Sidanius, J., Stallworth, L. M., & Malle, B. F. (1994). Social dominance orientation: A personality variable predicting social and political attitudes. Journal of personality and social psychology, 67(4), 741.

59. Sellaro, R., Derks, B., Nitsche, M. A., Hommel, B., van den Wildenberg, W. P., van Dam, K., & Colzato, L. S. Reducing prejudice through brain stimulation. Brain stimulation, 8(5), 891–897 (2015).

60. Liao, C., Wu, S., Luo, Y. J., Guan, Q., & Cui, F. Transcranial direct current stimulation of the medial prefrontal cortex modulates the propensity to help in costly helping behavior. Neuroscience Letters, 674, 54–59 (2018).

61. Huang, Y., Datta, A., Bikson, M., & Parra, L. C. ROAST: an open-source, fully-automated, realistic volumetric-approach-based simulator for TES. In 2018 40th Annual International Conference of the IEEE Engineering in Medicine and Biology Society (EMBC) (pp. 3072–3075). IEEE (2018).

62. Huang, Y., Datta, A., Bikson, M., & Parra, L. C. Realistic volumetric-approach to simulate transcranial electric stimulation—ROAST—a fully automated open-source pipeline. Journal of neural engineering, 16(5), 056006 (2019).

