## Supplementary file for "Causal Role of the Medial Prefrontal Cortex in Learning Social Hierarchy"

**This Supplementary file includes:**

Supplementary Information Text

Figures S1

Tables S1 to S6

**Supplementary Information Text**

***Supplementary analysis of the first block learning effect***

To justify that the brain stimulation effect was not due to the initial learning value within the group, we performed an additional analysis of the first block of the training phase. Note that within each training block participant repeated the learning of adjacent items twice. Thus, the average accuracy of the first block actually already included the learning effect through trial and errors. To address the question of whether there is a different initial value between groups, we conducted two analyses: 1) Repeated measure ANOVA with Hierarchy Type (Social, Non-social) * tDCS (Cathode, Sham, Anode) * Familiarity (First-time trials, Second-time trials) on the first block performance accuracy (%) and reaction time (ms); 2) One-sample Student t-test of performance accuracy of each condition to test of the average accuracy is significantly greater than 0.5 chance level.

As shown in Figure S1 A-B, the results of accuracy showed only a significant main effect of familiarity between the first-time and second-time trials ($F_{(1,125)}$ = 68.956, $P$ < 0.001, $\eta_p^2$ = 0.356; First-time trials: M=0.398, SD=0.196; Second-time trials: M=0.538, SD=0.204), indicating the participants in all the groups performed better in the second-time trials compared to the first-time, and thus there was a learning effect within the first block. However, there was no significant of three-way interaction ($F_{(2,125)}$ = 1.014, $P$ = 0.366, $\eta_p^2$ = 0.016) or two-way interaction between either hierarchy type and tDCS groups ($F_{(2,125)}$ = 1.014, $P$ = 0.366, $\eta_p^2$ = 0.016), or Familiarity and tDCS groups ($F_{(2,125)}$ = 2.606, $P$ = 0.078, $\eta_p^2$ = 0.040). These results indicate that for first-time and second-time trials, we did not observe differences between groups but only a general increased accuracy for the second-time trials within the first block. This provides evidence that there was no significant difference in the initial level of performance between the groups.

The analysis of reaction time showed similar results (see Figure S1 C-D), i.e., within groups participants responded significantly slower during second-time trials compared to the first-time ($F_{(1,125)}$ = 68.956, $P$ < 0.05, $\eta_p^2$ = 0.046; First-time trials: M=1501.073, SD=356.07; Second-time trials: M=1545.291, SD=337.691). They also responded faster in the non-social condition compared to the social hierarchy training ($F_{(1,125)}$ = 15.978, $P$ < 0.001, $\eta_p^2$ = 0.113; Non-social: M=1471.743, SD=348.474; Social: M=1574.621, SD=339.199). However, there was no significant three-way interaction ($F_{(2,125)}$ = 0.589, $P$ = 0.556, $\eta_p^2$ = 0.009) or two-way interaction in either hierarchy type and tDCS groups ($F_{(2,125)}$ = 0.089, $P$ = 0.915, $\eta_p^2$ = 0.001), or Familiarity and tDCS groups ($F_{(2,125)}$ = 1.223, $P$ = 0.298, $\eta_p^2$ = 0.019). These results further confirm that at the beginning of the task, there was no group difference but only a general learning effect such that on the second-time trails, after the feedback from first-time trails, participants spend more time to make a judgement and, compared to the non-social hierarchy, they took more time to make these judgements in the social condition. Finally, a second one-sample student t-test (>0.5 chance level, see table S1) showed that only the Anode groups had significantly higher than chance levels of accuracy on the second-time trials of the non-social hierarchy training. The other groups (cathode and sham) and all groups for the social condition were not significantly better than chance level, indicating that participants were still at an initial phase of learning the hierarchies.

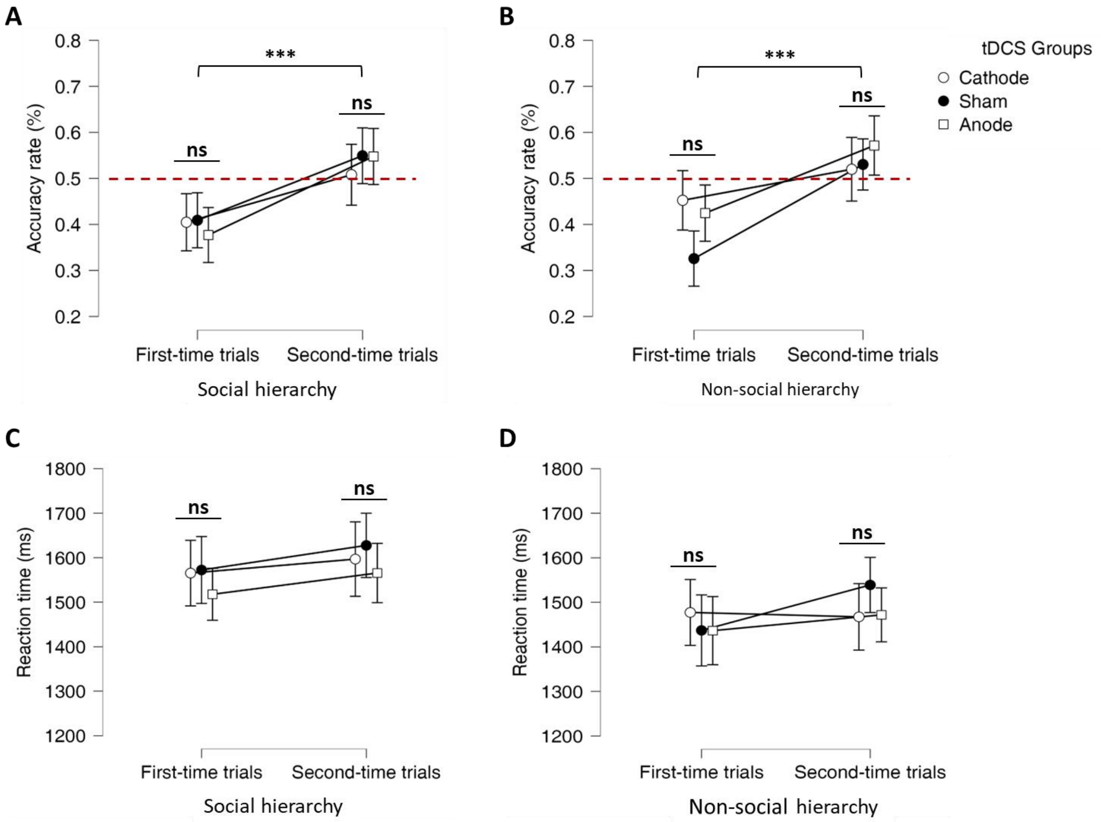


**Figure S1 First block analysis of training phase A) Social hierarchy and B) Non-social hierarchy accuracy; C) Social and D) Non-social hierarchy reaction time.** (*indicates P<0.05, **indicates P<0.005, ***indicates P<0.001; ****indicates P<0.0001; ns indicates non-significant; Error bars show SEM)

**Table S1**

| | | | | | | | | 95% CI for Cohen's d | |
|---|---|---|---|---|---|---|---|---|---|
| **Hierarchy type** | **Familiarity** | **tDCS** | **Mean** | **SD** | **t** | **p** | **Cohen's d** | **Lower** | **Upper** |
| Non-social | First-time | Anode | 0.425 | 0.195 | -2.286 | 0.986 | -0.357 | -0.62 | ∞ |
| | | Cathode | 0.452 | 0.231 | -1.338 | 0.906 | -0.209 | -0.467 | ∞ |
| | | Sham | 0.326 | 0.183 | -6.312 | 1 | -0.952 | -1.247 | ∞ |
| | Second-time | Anode | 0.571 | 0.202 | 2.293 | 0.014* | 0.358 | 0.091 | ∞ |
| | | Cathode | 0.52 | 0.233 | 0.666 | 0.255 | 0.104 | -0.154 | ∞ |
| | | Sham | 0.53 | 0.184 | 1.091 | 0.141 | 0.164 | -0.086 | ∞ |
| Social | First-time | Anode | 0.377 | 0.188 | -4.107 | 1 | -0.641 | -0.92 | ∞ |
| | | Cathode | 0.405 | 0.188 | -3.147 | 0.998 | -0.492 | -0.761 | ∞ |
| | | Sham | 0.409 | 0.174 | -3.464 | 0.999 | -0.522 | -0.784 | ∞ |
| | Second-time | Anode | 0.548 | 0.206 | 1.379 | 0.088 | 0.215 | -0.046 | ∞ |
| | | Cathode | 0.508 | 0.198 | 0.26 | 0.398 | 0.041 | -0.217 | ∞ |
| | | Sham | 0.549 | 0.205 | 1.592 | 0.059 | 0.24 | -0.013 | ∞ |

Note. For the Student t-test, the alternative hypothesis specifies that the mean is greater than 0.5. *<0.05

**Table S2. Descriptive statistics of Demographic, Questionnaires, and Task-relevant subjective rating**

| | | ***Anodal (N=42)*** | ***Sham (N=44)*** | ***Cathodal (N=42)*** |
|---|---|---|---|---|
| ***Gender*** | | Males=21<br>Females=21 | Males=22<br>Females=23 | Males=21<br>Females=21 |
| ***Age*** | | 19.610±0.231 | 19.909±0.239 | 20.171±0.281 |
| ***SDO*** | | 51.190±13.515 | 53.114±12.085 | 51.595±10.378 |
| ***Truth Degree*** | | 5.452±0.350 | 5.545±0.337 | 5.976±0.352 |
| ***tDCS uncomfortable rating*** | | 1.536±0.146 | 1.341±0.101 | 1.762±0.137 |
| ***Choice Bias*** | **Training** | Left=0.501±0.005<br>Right=0.499±0.005 | Left=0.504±0.005<br>Right=0.496±0.005 | Left=0.503±0.005<br>Right=0.497±0.005 |
| | **Test** | Left=0.502±0.003<br>Right=0.498±0.003 | Left=0.508±0.003<br>Right=0.492±0.003 | Left=0.499±0.003<br>Right=0.501±0.003 |

Note: One-way ANOVA showed no significant differences of the above measurements among groups.

**Table S3. tDCS effect of block-to-block hierarchy learning in the performance % of Social condition**

| | ***Anode vs Sham*** | | | | ***Cathode vs Sham*** | | | |
|---|---|---|---|---|---|---|---|---|
| | **Training** | | **Test** | | **Training** | | **Test** | |
| ***Block*** | **b(SE)** | **P** | **b(SE)** | **P** | **b(SE)** | **P** | **b(SE)** | **P** |
| ***1*** | -0.053(0.026) | 0.044* | -0.113(0.036) | 0.002** | 0.002(0.026) | 0.925 | 0.037(0.036) | 0.305 |
| ***2*** | -0.055(0.025) | 0.028* | -0.103(0.035) | 0.004** | 0.009(0.025) | 0.728 | 0.026(0.035) | 0.451 |
| ***3*** | -0.055(0.024) | 0.020* | -0.090(0.035) | 0.009** | 0.014(0.023) | 0.554 | 0.016(0.033) | 0.624 |
| ***4*** | -0.055(0.023) | 0.016* | -0.076(0.033) | 0.022* | 0.018(0.022) | 0.418 | 0.008(0.032) | 0.803 |
| ***5*** | -0.053(0.022) | 0.014* | -0.063(0.031) | 0.045* | 0.021(0.021) | 0.320 | 0.001(0.030) | 0.971 |
| ***6*** | -0.050(0.021) | 0.015* | -0.051(0.029) | 0.082 | 0.022(0.019) | 0.254 | -0.004(0.028) | 0.882 |
| ***7*** | -0.046(0.019) | 0.016* | -0.040(0.026) | 0.133 | 0.023(0.018) | 0.209 | -0.008(0.025) | 0.761 |
| ***8*** | -0.042(0.018) | 0.019* | -0.030(0.023) | 0.197 | 0.022(0.016) | 0.180 | -0.010(0.023) | 0.662 |
| ***9*** | -0.038(0.017) | 0.022* | -0.023(0.021) | 0.268 | 0.021(0.015) | 0.160 | -0.011(0.021) | 0.583 |
| ***10*** | -0.033(0.015) | 0.027* | -0.017(0.018) | 0.344 | 0.019(0.013) | 0.146 | -0.012(0.018) | 0.520 |
| ***11*** | -0.029(0.014) | 0.032* | -0.012(0.015) | 0.420 | 0.017(0.012) | 0.136 | -0.011(0.016) | 0.470 |
| ***12*** | -0.025(0.012) | 0.038* | -0.009(0.013) | 0.495 | 0.016(0.010) | 0.129 | -0.011(0.014) | 0.429 |

Significance: $^{*}p < 0.05$, $^{**}p < 0.01$, $^{***}p < 0.001$.

**Table S4. tDCS effect of block-to-block hierarchy learning in the reaction time of Social condition**

| | *Anode vs Sham* | | | | *Cathode vs Sham* | | | |
|---|---|---|---|---|---|---|---|---|
| | **Training** | | **Test** | | **Training** | | **Test** | |
| ***Block*** | **b(SE)** | **P** | **b(SE)** | **P** | **b(SE)** | **P** | **b(SE)** | **P** |
| ***1*** | -32.440(37.704) | 0.390 | 35.783(41.108) | 0.384 | -27.055(37.704) | 0.473 | 2.289(41.108) | 0.956 |
| ***2*** | -22.860(37.052) | 0.537 | 45.753(40.126) | 0.254 | -26.770(37.052) | 0.470 | 3.080(40.126) | 0.939 |
| ***3*** | -13.280(36.522) | 0.716 | 55.723(39.323) | 0.156 | -26.485(36.522) | 0.468 | 3.871(39.323) | 0.922 |
| ***4*** | -3.7008(36.120) | 0.918 | 65.693(38.710) | 0.090 | -26.200(36.120) | 0.468 | 4.663(38.710) | 0.904 |
| ***5*** | 5.879(35.849) | 0.870 | 75.663(38.296) | 0.048* | -25.915(35.849) | 0.470 | 5.454(38.296) | 0.887 |
| ***6*** | 15.459(35.713) | 0.665 | 85.633(38.087) | 0.025* | -25.630(35.713) | 0.473 | 6.245(38.087) | 0.870 |
| ***7*** | 25.038(35.713) | 0.483 | 95.604(38.087) | 0.012* | -25.345(35.713) | 0.478 | 7.037(38.087) | 0.853 |
| ***8*** | 34.618(35.849) | 0.334 | 105.574(38.296) | 0.006** | -25.060(35.849) | 0.485 | 7.828(38.296) | 0.838 |
| ***9*** | 44.198(36.120) | 0.221 | 115.544(38.710) | 0.003** | -24.775(36.120) | 0.493 | 8.619(38.710) | 0.824 |
| ***10*** | 53.777(36.522) | 0.141 | 125.514(39.323) | 0.001** | -24.491(36.522) | 0.502 | 9.411(39.323) | 0.811 |
| ***11*** | 63.357(37.052) | 0.087 | 135.484(40.126) | 0.001** | -24.206(37.052) | 0.514 | 10.202(40.126) | 0.799 |
| ***12*** | 72.937(37.704) | 0.053 | 145.454(41.108) | <.0001 *** | -23.921(37.704) | 0.526 | 10.993(41.108) | 0.789 |

Significance: $^{*}p < 0.05$, $^{**}p < 0.01$, $^{***}p < 0.001$.

**Table S5. tDCS effect of block-to-block hierarchy learning in the % performance of Non-Social condition**

| | ***Anode vs Sham*** | | | | ***Cathode vs Sham*** | | | |
|---|---|---|---|---|---|---|---|---|
| | **Training** | | **Test** | | **Training** | | **Test** | |
| ***Block*** | **b(SE)** | **P** | **b(SE)** | **P** | **b(SE)** | **P** | **b(SE)** | **P** |
| ***1*** | 0.007(0.026) | 0.788 | -0.053(0.036) | 0.142 | 0.022(0.026) | 0.403 | -0.004(0.037) | 0.907 |
| ***2*** | 0.005(0.025) | 0.828 | -0.045(0.035) | 0.197 | 0.010(0.025) | 0.682 | -0.017(0.036) | 0.639 |
| ***3*** | 0.004(0.024) | 0.873 | -0.037(0.034) | 0.272 | -0.001(0.024) | 0.969 | -0.027(0.034) | 0.422 |
| ***4*** | 0.002(0.023) | 0.920 | -0.029(0.032) | 0.364 | -0.011(0.023) | 0.631 | -0.036(0.033) | 0.268 |
| ***5*** | 0.001(0.022) | 0.966 | -0.022(0.031) | 0.468 | -0.019(0.022) | 0.371 | -0.043(0.031) | 0.168 |
| ***6*** | 0.000(0.021) | 0.992 | -0.016(0.029) | 0.575 | -0.026(0.021) | 0.207 | -0.048(0.030) | 0.107 |
| ***7*** | -0.001(0.019) | 0.955 | -0.011(0.026) | 0.680 | -0.031(0.020) | 0.115 | -0.050(0.028) | 0.071 |
| ***8*** | -0.002(0.018) | 0.923 | -0.007(0.024) | 0.778 | -0.034(0.018) | 0.066 | -0.052(0.026) | 0.049* |
| ***9*** | -0.002(0.016) | 0.896 | -0.004(0.022) | 0.867 | -0.035(0.017) | 0.040* | -0.052(0.025) | 0.035* |
| ***10*** | -0.002(0.015) | 0.873 | -0.001(0.02) | 0.947 | -0.035(0.016) | 0.026* | -0.051(0.023) | 0.027* |
| ***11*** | -0.002(0.013) | 0.854 | 0.000(0.017) | 0.983 | -0.035(0.015) | 0.018* | -0.049(0.021) | 0.021* |
| ***12*** | -0.002(0.012) | 0.838 | 0.002(0.015) | 0.921 | -0.033(0.013) | 0.013* | -0.046(0.020) | 0.018* |

Significance: $^{*}p < 0.05$, $^{**}p < 0.01$, $^{***}p < 0.001$.

**Table S6. tDCS effect of block-to-block hierarchy learning in the reaction time of Non-Social condition**

| | ***Anode vs Sham*** | | | | ***Cathode vs Sham*** | | | |
|---|---|---|---|---|---|---|---|---|
| | **Training** | | **Test** | | **Training** | | **Test** | |
| ***Block*** | **b(SE)** | **P** | **b(SE)** | **P** | **b(SE)** | **P** | **b(SE)** | **P** |
| *1* | -79.414(37.704) | 0.035* | -87.487(41.108) | 0.033* | -62.531(37.704) | 0.097 | -93.396(41.108) | 0.023* |
| *2* | -71.176(37.052) | 0.055 | -73.214(40.126) | 0.068 | -53.684(37.052) | 0.147 | -74.721(40.126) | 0.063 |
| *3* | -62.937(36.522) | 0.085 | -58.940(39.323) | 0.134 | -44.837(36.522) | 0.220 | -56.046(39.323) | 0.154 |
| *4* | -54.698(36.120) | 0.130 | -44.667(38.710) | 0.249 | -35.990(36.120) | 0.319 | -37.371(38.710) | 0.334 |
| *5* | -46.460(35.849) | 0.195 | -30.394(38.296) | 0.427 | -27.143(35.849) | 0.449 | -18.695(38.296) | 0.625 |
| *6* | -38.221(35.713) | 0.285 | -16.121(38.087) | 0.672 | -18.296(35.713) | 0.608 | -0.020(38.087) | 1.000 |
| *7* | -29.982(35.713) | 0.401 | -1.847(38.087) | 0.961 | -9.449(35.713) | 0.791 | 18.655(38.087) | 0.624 |
| *8* | -21.744(35.849) | 0.544 | 12.426(38.296) | 0.746 | -0.602(35.849) | 0.987 | 37.330(38.296) | 0.330 |
| *9* | -13.505(36.120) | 0.708 | 26.699(38.710) | 0.490 | 8.245(36.120) | 0.819 | 56.005(38.71) | 0.148 |
| *10* | -5.267(36.522) | 0.885 | 40.972(39.323) | 0.297 | 17.092(36.522) | 0.640 | 74.680(39.323) | 0.058 |
| *11* | 2.972(37.052) | 0.936 | 55.246(40.126) | 0.169 | 25.938(37.052) | 0.484 | 93.355(40.126) | 0.020* |
| *12* | 11.211(37.704) | 0.766 | 69.519(41.108) | 0.091 | 34.785(37.704) | 0.356 | 112.03(41.108) | 0.006** |

Significance: $^{*}p < 0.05$, $^{**}p < 0.01$, $^{***}p < 0.001$.
